# PP-2, a src-kinase inhibitor, is a potential corrector for F508del-CFTR in cystic fibrosis

**DOI:** 10.1101/288324

**Authors:** Yunguan Wang, Kavisha Arora, Fanmuyi Yang, Woong-Hee Shin, Jing Chen, Daisuke Kihara, Anjaparavanda P. Naren, Anil G. Jegga

## Abstract

Cystic fibrosis (CF) is an autosomal recessive disorder caused by mutations in the CF transmembrane conductance regulator (CFTR) gene. The most common mutation in CF, an in-frame deletion of phenylalanine 508, leads to a trafficking defect and endoplasmic reticulum retention of the protein where it becomes targeted for degradation. Successful clinical deployments of ivacaftor and ivacaftor/lumacaftor combination have been an exciting translational development in treating CF. However, their therapeutic effects are variable between subjects and remain insufficient. We used the Library of Integrated Network-based Cellular Signatures (LINCS) database as our chemical pool to screen for candidates. For *in silico* screening, we integrated connectivity mapping and CF systems biology to identify candidate therapeutic compounds for CF. Following *in silico* screening, we validated our candidate compounds with (i) an enteroid-based compound screening assay using CF (ΔF508/ΔF508-CFTR) patient-derived enteroids, (ii) short-circuit current analysis using polarized CF primary human airway epithelial cells and (iii) Western blots to measure F508-del-CFTR protein maturation. We identified 184 candidate compounds with *in silico* screening and tested 24 of them with enteroid-based forskolin-induced swelling (FIS) assay. The top hit compound was PP2, a known src-kinase inhibitor that induced swelling in enteroid comparable to known CF corrector (lumacaftor). Further validation with Western blot and short-circuit current analysis showed that PP-2 could correct mutant CFTR mis-folding and restore CFTR-mediated transmembrane current. We have identified PP2, a known src-kinase inhibitor, as a novel corrector of ΔF508-CFTR. Based on our studies and previous reports, src kinase inhibition may represent a novel paradigm of multi-action therapeutics – corrector, anti-inflammatory, and anti-infective – in CF.

## Background

Cystic fibrosis (CF) is a life-limiting genetic disorder affecting 70,000 individuals worldwide (about 30,000 in the United States). Although the recent drug approvals for CF (potentiator ivacaftor alone and in combination with corrector lumacaftor) are promising, the pursuit for additional therapeutics for CF needs to be continued [1] for the following reasons. *In vitro* studies have shown that the approved combinatorial has limited clinical efficacy with potential interference of potentiator (augment CFTR channel function) with corrector (promote the read-through of nonsense mutations or facilitate the translation, folding, maturation, and trafficking of mutant CFTR to the cell surface) actions and destabilization of corrected ΔF508-CFTR, the most common mutant in CF [2-4]. Additionally, apart from being cost-prohibitive (>$250K/year), the approved combinatorial will not work for all CF patients [2, 3, 5, 6]. The context of various CFTR mutations (>2000), the complexity of the underlying pathways, and pathway cross-talk suggest that a computational big data approach that uses high-throughput experimental data and systems biology attributes could lead to the discovery of hitherto unanticipated potential targets and therapeutic candidates for CF. Indeed, mining some of the genomics and small-molecule data silos (e.g., disease/drug gene expression profiles or signatures) in an integrative manner has already been shown to be of translational benefit such as finding novel drug–drug and drug– disease relationships [7-17]. However, a majority of such signature comparison-based approaches treats cells as black boxes; gene expression patterns from small-molecule treatments and from patients or disease models are the inputs and a ranked list of compounds is the output. Further, the output from such computational analysis is often large, necessitating further prioritization. Thus, there is a critical and unmet need to triage small molecules discovered from high-throughput screens for further development so that investigators can focus on a smaller number with greater success and lower cost. By incorporating additional layers of systems biology attributes (e.g., wild type and mutant CFTR-specific protein interactions and CF-relevant signaling networks and pathways), the current study sought to unveil the internal configuration of the black box underlying the connected small molecules, their targets, and CF disease gene signatures. Such understanding could lead to identifying new therapeutics, drug targets, and target pathways and novel mechanisms of action for CF. Additionally, doing these studies earlier in the drug discovery pipeline could help in potentially foreseeing or avoiding late-stage clinical trial failures. Since the compound screening framework in the current study includes approved drugs, drug repositioning candidates for CF might also be found.

The CF gene expression signature derived from rectal epithelia (RE) of human CF patients with ΔF508-CFTR mutation [18] was used to search the Library of Integrated Network-based Cellular Signatures (LINCS) database [19, 20] to identify compounds that are anti-correlated with the CF signature. We then complemented this unbiased chemical signature-based screening method with CF knowledge-based systems biology approaches to characterize and infer mechanistic insights. Finally, we tested computationally prioritized small molecules from this integrated approach *in vitro* for their effect on fluid secretion (in the presence and absence of CF-corrector VX809) by using intestinal organoids (enteroids or miniguts) generated from the intestines of ΔF508-CFTR homozygous mice and CF patients. Additional candidate compounds were validated by measuring the CFTR-dependent short circuit currents (*I*_sc_) by using polarized CF primary human airway epithelial cells [21, 22].

## Methods

### Differentially expressed genes in CF

Microarray data of rectal epithelial cells from healthy volunteers and from CF patients with ΔF508-CFTR mutation (homozygous) were downloaded from the National Center for Biotechnology Information Gene Expression Omnibus (GEO) [23]. Differential analysis for genes was performed using the R package limma with *P*=0.05 and fold change threshold at 1.5 [24].

### Computational small-molecule compound screening

We used the LINCS cloud web tool [20] and gene set enrichment analysis (GSEA) to identify small molecules from the NIH’s LINCS library that could potentially reverse the CF gene expression profile. LINCS signatures (460,000) were downloaded from LINCScloud (www.lincscloud.org) using API. Each signature consisted of a list of 100 most up-regulated and 100 most down-regulated probes. The original ConnectivityMap method [14] was applied to a CF disease signature against each of the 460,000 LINCS signatures to calculate a connectivity score. This resulted in 460,000 LINCS Connectivity Scores for each CF disease signature. Since in LINCS experiment the same treatment was applied in multiple cell lines with different duration and dosages, they were all considered biological replicates of the same treatment in the current analysis. Therefore, a non-parametric two sample Kolmogorov–Smirnov test was used to compare the Connectivity Scores of signatures from one treatment against the background Connectivity Scores of the entire 460,000 LINCS signatures. Since we were interested in the compound whose signature was inversely correlated to the CF disease signature, the one-tailed KS test was applied wherein the alternative hypothesis was that the Connectivity Scores of the underlying treatment were higher than the background Connectivity Scores (note that negative Connectivity Score indicates reverse relation between the treatment and CF disease signature). This one-tailed two sample KS test was performed for each unique compound treatment, and the resulting p-value was reported as the significance of the compound. This p-value was then used to rank all compound treatments in LINCS. The complete process was performed using R environment. LINCS signatures were downloaded using “jsonlite” package [25].

### Computational modeling of ΔF508-CFTR and docking of PP2 and VX-809

Open-state WT CFTR structure was obtained from a previous work by Dalton *et al.* [26]. ΔF508-CFTR mutant structure was predicted by Modeller9v18 [27] using wild type structure as a template. Three-dimensional ligand structures of VX-809 and PP2 were obtained from PubChem [28], and adding hydrogens to the receptor and assigning Gasteiger charges to the protein and the compounds were done by AutoDockTools [29].

### Reagents and antibodies

PP2 and other compounds tested were purchased from Tocris Bioscience (Ellisville, Missouri) while VX-809 and VX-661 were obtained from Selleck (Houston, Texas).

### Human studies

Human intestinal biopsy and lung tissues were obtained from CFTRWT/WT non-CF and ΔF508/ΔF508-CFTR CF individuals under the protocol and consent form approved by the Institutional Review Board at the Cincinnati Children’s Hospital Medical Center (IRB # 2011-2616). All adult participants provided informed consent, and a parent or guardian of any child participant provided informed permission on the child’s behalf. All consent was obtained in written form.

### Enteroid cultures

Mouse intestinal organoids were cultured as described previously (25). Human duodenal crypt isolation and enteroid expansion was performed as described previously (https://www.jove.com/video/52483/establishment-human-epithelial-enteroids-colonoids-from-whole-tissue) with some adaptations. Briefly, fresh biopsy is rinsed in ice-cold Dulbecco’s Phosphate buffered saline without Ca^2+^ and Mg^2+^ (DPBS, Gibco), mounted and immersed in DPBS in a silica gel coated petri-dish using minutien pins with the mucosal side facing up.

Mucosa is gently scraped with curved forceps to remove villi and debris followed by 3-4 washes with DPBS. Crypts were dissociated using 2 mM EDTA (30 min, 4°C with gentle shaking) followed by gentle scraping of the mucosa. The crypt suspension was filtered through a 150 µm nylon mesh twice and pelleted at 50 × g, 4°C. The crypt pellet was resuspended in matrigel matrix (200 to 500 crypts/50 µl matrigel per well of a 24 well plate). Matrigel was allowed to polymerize by placing the plate in a 37°C, 5% CO2 incubator for 30 min followed by addition of complete growth factor supplemented human minigut medium (Advanced DMEM/F12 medium with 2 mM glutamine, 10 mM HEPES, 100 U/mL penicillin, 100 g/mL streptomycin, 1 N2 supplement, 1 B27 supplement and 1% BSA supplemented with 50% Wnt-3A-conditioned medium, 1 µg/ml R-spondin 1, 100 ng/ml Noggin. 50 ng/mL EGF, 500 nM A-83-01, 10 µM SB202190, 10 nM [Leu]15-Gastrin 1, 10 mM Nicotinamide and 1 mM N-Acetylcysteine.

### Fluid secretion measurement in intestinal spheroids

Isolation of intestinal spheres and measurement of fluid secretion were performed as described previously [30]. Day 1-4 intestinal organoids were treated with 0.1-10 µM of the test compound for 24 h before stimulation of CFTR function using forskolin (10 µM). Fluid secretion measurements were done before and after 30 min of a stimulation period for mouse organoids and 120 min for human organoids. Quantitation of fluid secretion in the intestinal spheres was performed as described previously [30, 31].

### I_sc_ measurement

Primary human ΔF508/ΔF508 *CFTR* bronchial epithelial cells grown on Costar Transwell permeable supports (Cambridge, MA; filter diameter 12 mm) were mounted in an Ussing chamber maintained at 37°C. Epithelial cells were pretreated with DMSO, PP2 (2 µM, 24 h), VX-809 (2 µM, 24 h), and PP2 + VX-809. A 2-mV pulse was applied every 1 min throughout the experiment to check the integrity of the epithelia. Cells were treated with 50 µM amiloride at the beginning of the experiment. After stabilizing *I_sc_*, cells were treated with forskolin (Fsk) (10 µM) on the apical side. CFTR_inh_-172 (20-50 μM) was added to the apical side at the end of each experiment to verify current dependence on CFTR.

### PP2 RNA-Seq

RNA sequencing was performed on primary human bronchial epithelial cells homozygous for ΔF508-CFTR mutation (sample ID KKCFFT004I, Charles River, Wilmington, MA). Briefly, fully differentiated bronchial epithelial cells maintained on air-liquid interface were treated with DMSO or PP2 (2 µM, 24 h). Total RNA was isolated using mirVana™ miRNA Isolation Kit (Carlsbad, California) and used for RNA sequencing.

## Results

### Identifying candidate anti-CF small molecules through integrated gene expression profiling and systems biology

We used a published CF patient gene expression data set (GSE15568; [18], which was based on rectal epithelia samples of CF patients with ΔF508-CFTR mutation. Differential analysis of gene expression was performed by the R package ‘limma’ with P-value threshold at 0.05 and fold change threshold at 1.5 [24]. Compared with healthy controls, 1330 genes were differentially expressed in ΔF508-CF patients. Among these genes, 555 were downregulated, and 775 were upregulated (Additional file 1). We used this differentially expressed gene (DEG) set to identify small molecules from the NIH’s LINCS library that could potentially reverse the CF gene expression profile. We used the LINCS cloud web tool and gene set enrichment analysis (GSEA) for this purpose (Fig. 1). Following this signature reversal step, 184 compounds were identified as significantly reversing the CF disease signature.

**Fig. 1.**
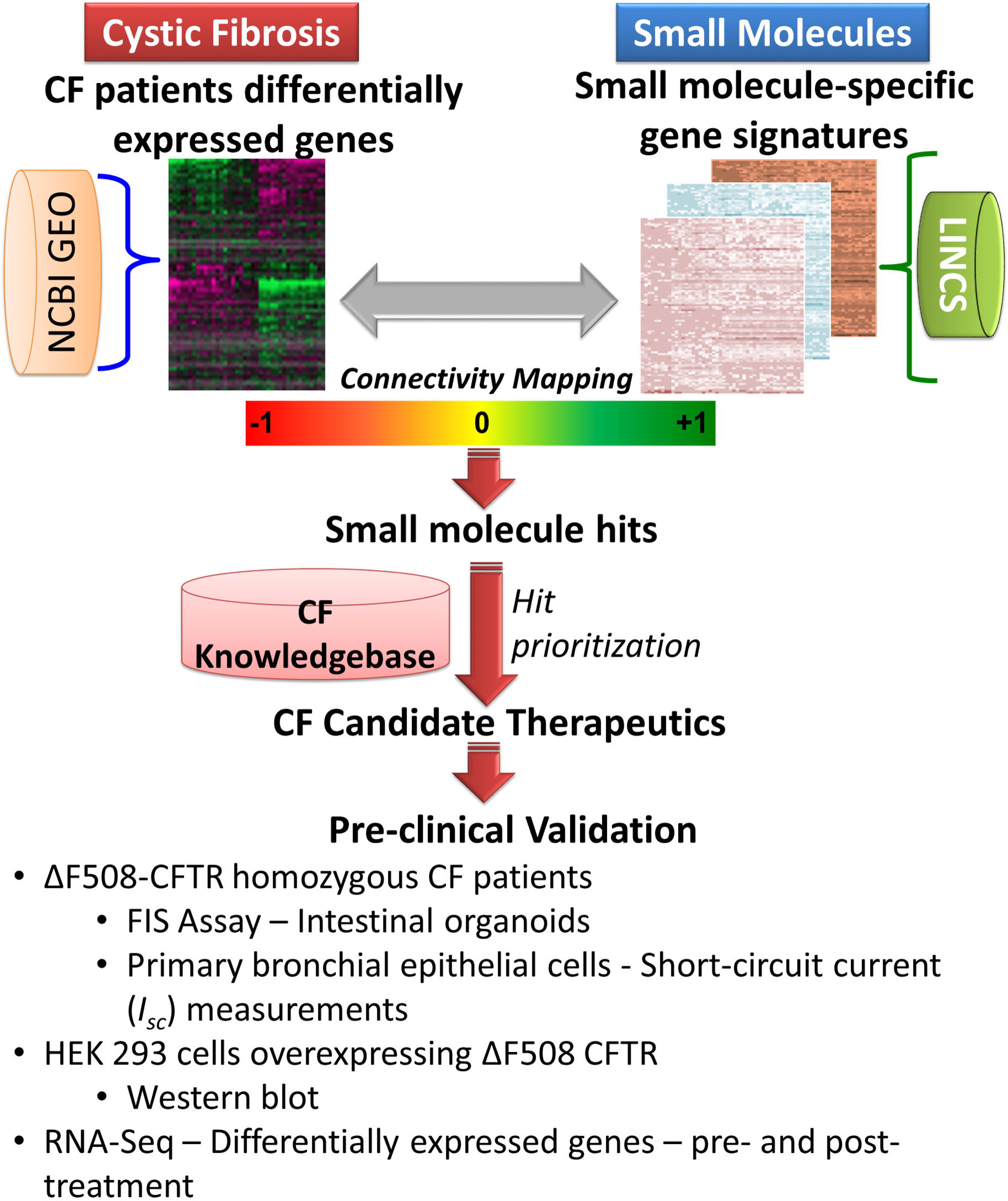
Schematic representation of workflow to identify CF correctors. Differentially expressed genes from CF patients (ΔF508-CFTR homozygous) were used for connectivity mapping to identify CF candidate therapeutics. These candidate therapeutics were further prioritized by incorporating additional layers of information from systems biology of CF. Prioritized CF candidate therapeutics were experimentally validated using intestinal organoids and bronchial epithelial cells from CF patients.

To elucidate computationally each of the 184 compounds, specifically to determine their putative functional relatedness to CFTR protein (WT or ΔF508) interactome [32] and to compiled CF-relevant pathways (Additional file 2, see Methods), we performed a singular enrichment analysis (SEA) by using the DEGs of each of these compounds in the A549 cell lines available from the LINCS database. Eight compounds that were not enriched in any of the CF-relevant pathways were removed from the candidate list of compounds. To further narrow the candidates potentially involved in CF, we selected 10 pathways that were highly related with CF pathogenesis and ranked all the remaining compounds based on their average enrichment p-values in these terms. To diversify our selection, we also selected a few compounds with low ranking scores. Finally, we randomly selected 18 small molecules from the top 1/3 ranked, 5 from the middle, and 1 from the bottom ranked compounds (Additional file 3). These compounds also represented different chemical classes (such as flavonoids, src family kinase inhibitors, MAPK inhibitors, mTOR inhibitor, and PI3K inhibitor; Additional file 4).

### Screening of candidate compounds in mouse ΔF508/ΔF508-*cft*r enteroids for mutant *CFTR* functional rescue

In order to screen for the effect of 24 selected candidates from the integrated computational and CF systems biology on ΔF508-CFTR functional rescue, we sorted to physiologically relevant intestinal stem cell-derived spheroids (enteroids) isolated from ΔF508/ΔF508-*cftr* mice. Intestinal spheroids are the validated models for studying CFTR-dependent fluid secretion [30, 31]. Of the 24 total compounds screened, two compounds (PP2 and LY-294002) demonstrated increased forskolin-induced swelling (FIS) in the enteroids compared to the DMSO-treated control (Fig. 2; Additional file 4). Among these, small-molecule PP2 showed the maximal rescue of ΔF508 *cftr* function, and the effects were noted to be higher than that with approved and investigational CF correctors (VX-809 and VX-661) 12% and 14%, respectively. We also observed a dose-dependent (0.1 μm to 10 μm) rescue of mutant CFTR in the presence of PP2 in the enteroids, and this effect was mitigated in the presence of CFTR_inh_-172 (Figs. 3A and 3B). A potential synergism was noted between PP2 and CF correctors VX-809 and VX-661 at 1 μm PP2 (Fig. 3B).

**Fig. 2.**
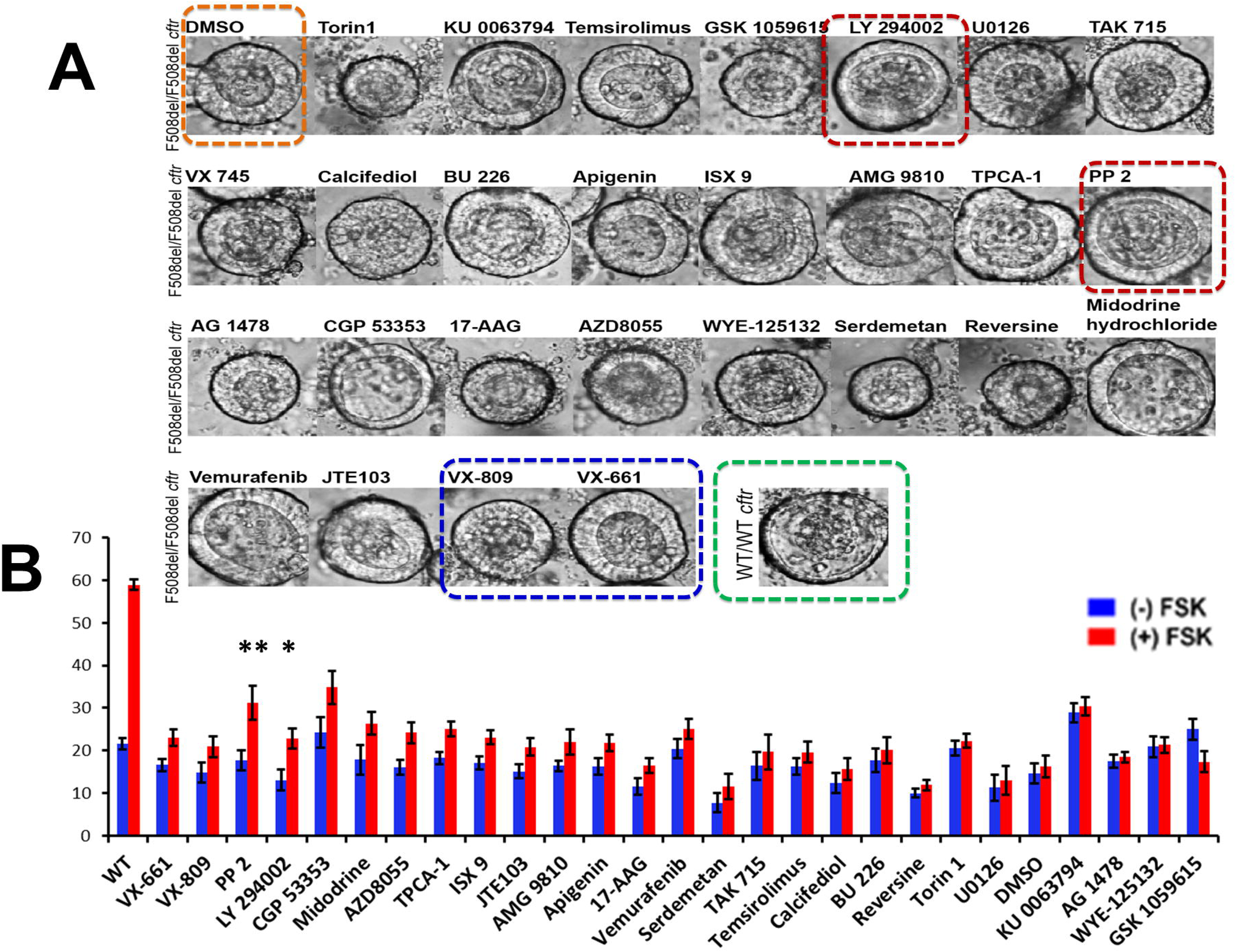
FIS in ΔF508/ΔF508-cftr intestinal organoids. Representative images of intestinal spheres demonstrating fluid secretion in response to the test compounds (Panel A). Bar graph represents quantitation of fluid secretion in the intestinal organoids (Panel B). In this preliminary screening assay, all compounds were tested at a dose of 10 µM, while the known correctors (VX-809 and VX-661) were tested at a dose of 2 µM. The highlighted compounds (PP2 and LY294002) in the bar graph were found to show forskolin-induced swelling significantly better than that seen with the treatment of VX-809 or VX-661. The related raw, and processed data for FIS assay is made available as Additional file 4.

**Fig. 3.**
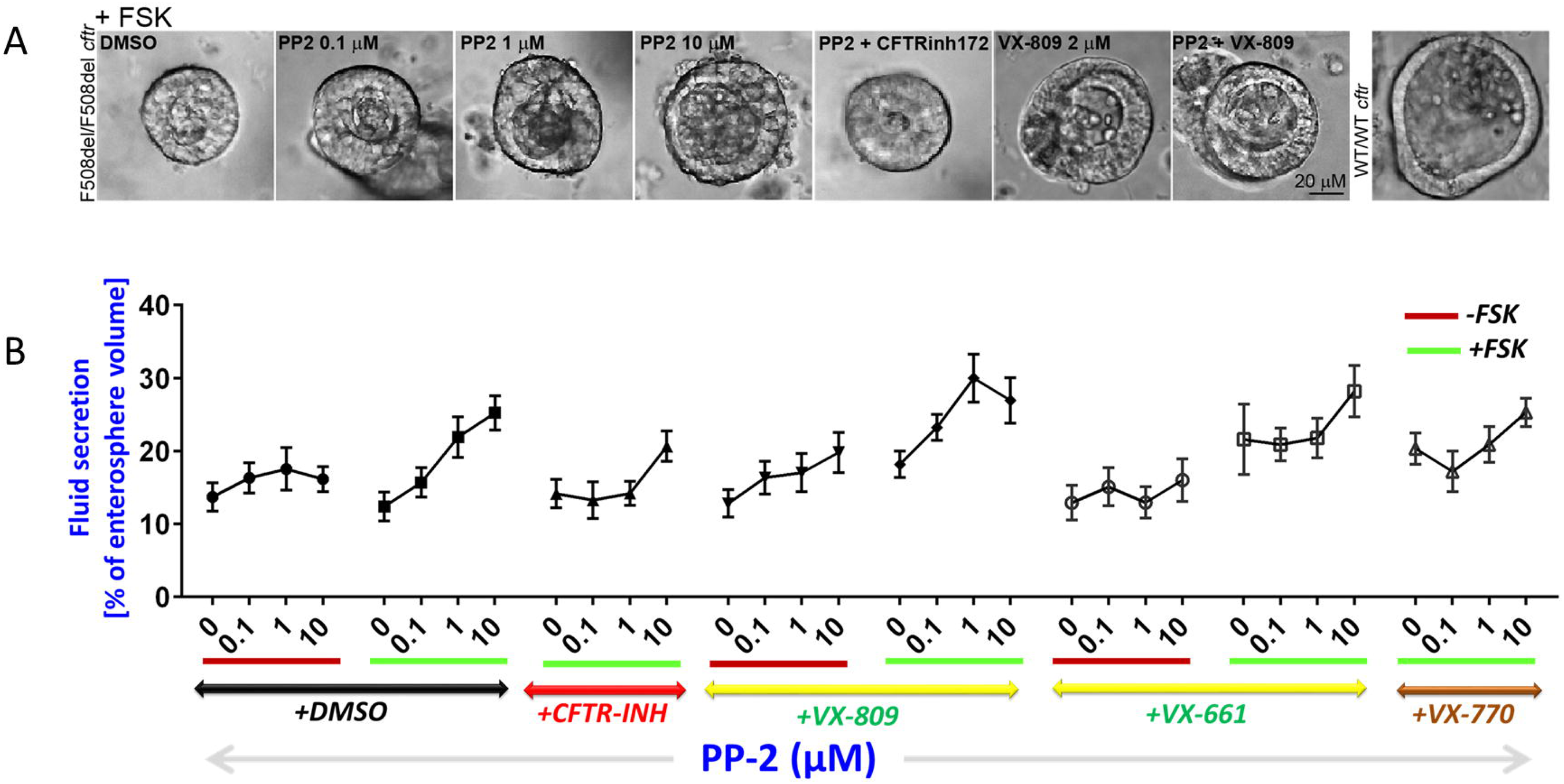
PP2 Dose response experiments in mice. **Panel A.** Representative images of ΔF508/ΔF508-*cftr* enterospheres depict secretion under various treatment conditions: (i) PP2 (0, 0.1, 1 and 10 µM, 24 h), (ii) PP2 (1 µM, 24 h) + CFTR_inh_-172 (20 µM, 30 min), (iii) VX-809 (2 µM, 24 h), and (iv) PP2 (1 µM, 24 h) + VX-809 (2 µM, 24 h). **Panel B**. Line graph represents quantitation of fluid secretion in enterospheres under various treatment conditions as described previously (Panel A).

### Screening of PP2 and its non-src-kinase-inhibitor analog PP3 using enteroids derived from CF patients (***Δ***F508/***Δ***F508-CFTR)

The corrector potential of PP2 could be recapitulated in CF (ΔF508/ΔF508-*CFTR*) patient-derived duodenal organoids. PP2 induced significant FIS at doses of 2 µM, and the enteroid swelling induced by PP2 was comparable to that by VX-809. To examine the dependence of PP2-mediated FIS on src kinase inhibition, we included PP3, an inactive PP2 analog, in our CFTR assay; the analog failed to mediate FIS (Fig. 4A).

**Fig. 4.**
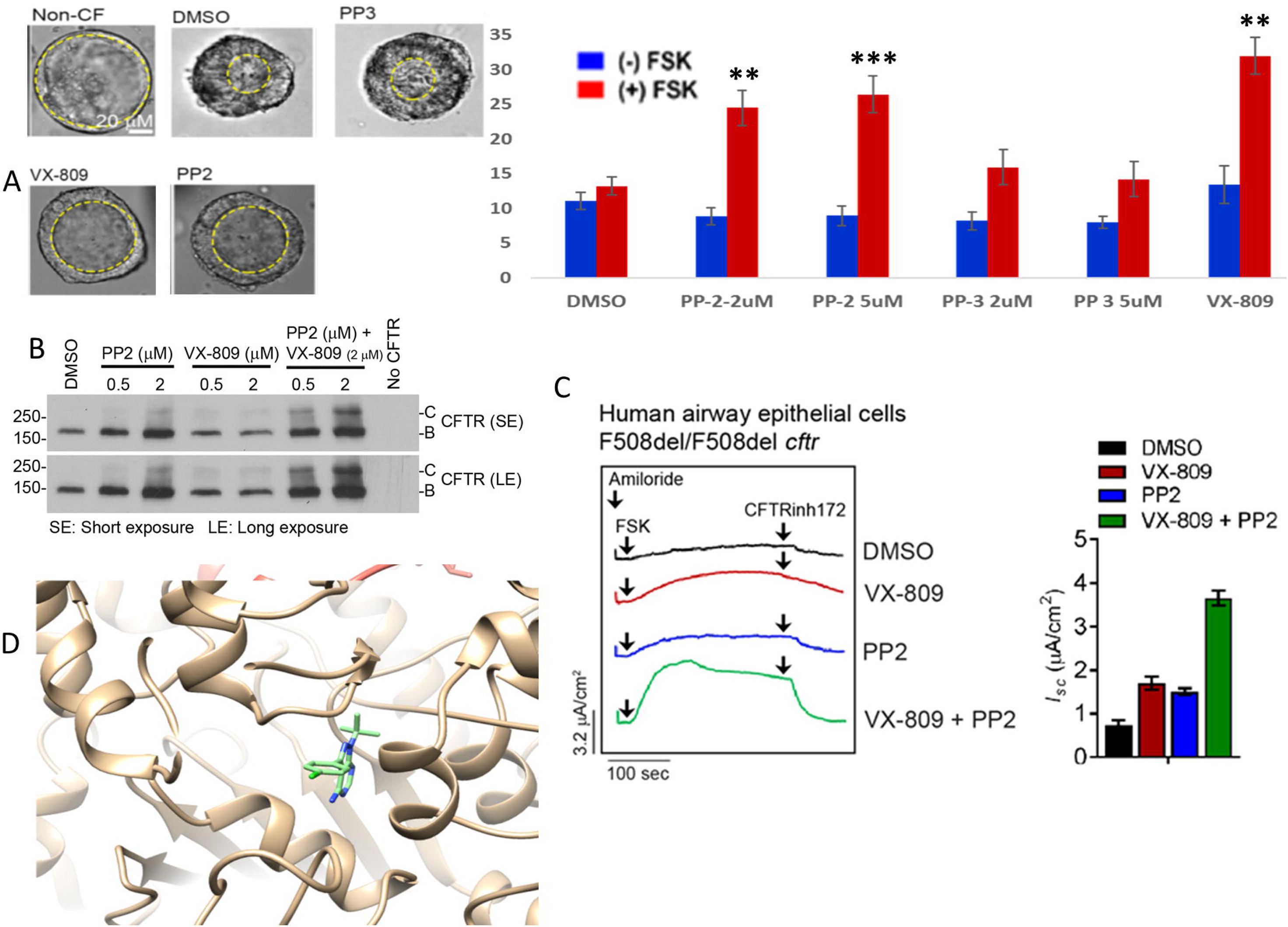
Preclinical validation of PP2. **Panel A** shows representative images of intestinal spheres demonstrating forskolin-induced swelling (FIS) in enteroids from normal subject and those from cystic fibrosis patients in response to DMSO, PP3, VX-809, and PP2. The bar graph represents quantitation of fluid secretion in the intestinal organoids. **Panel B.** Western blot data depict bands B (immature or endoplasmic reticulum form) and C (mature or membrane form) of CFTR immunoprecipitated from HEK 293 cells that overexpressed FLAG ΔF508-CFTR with and without treatment with PP2 (0.5 and 2 µM, 24 h), VX-809 (2 µM, 24 h) alone, and PP2 + VX-809 (2 µM each, 24 h). **Panel C**. Representative CFTR-mediated short-circuit currents (*I_sc_*) in human bronchial epithelial cells carrying ΔF508/ΔF50-CFTR in response to VX-809, PP2, and VX-809+PP2 treatment. Bar graphs represent data quantification of maximal increase in *I_sc_*/cm^2^ from n = 5 (DMSO-treated) and n=6 (PP2-treated) independent traces. **Panel D**. Predicted binding pose of PP2 to ATP binding site.

### PP2 corrected ***Δ***F508-*CFTR* mutant protein

Since we observed that PP2 functionally rescued ΔF508-CFTR, we sought to investigate whether PP2 rescues ΔF508-CFTR trafficking. Processing of immature band B of ΔF508-CFTR to mature band C would suggest improved trafficking or correction of the mutant ΔF508-CFTR protein. Treatment of ΔF508-CFTR expressing HEK 293 cells with PP2 increased the formation of mature band C, an effect like that of VX-809. Additionally, a strong synergistic effect was observed upon simultaneous treatment of cells with PP2 and VX-809 (Fig. 4B). This observation qualified PP2 as a candidate CF corrector. We validated the rescue potential of PP2 by measuring CFTR-mediated short circuit currents in primary human bronchial epithelial cells with ΔF508/ΔF508-CFTR mutation and observed a functional restoration of the mutant protein by ∼4-fold and synergistic functional rescue in the presence of PP2 and VX-809 combination (Fig. 4C).

### Computational modeling of ***Δ***F508-CFTR and PP2

To predict the PP2 mode of action (MoA), we docked PP2 and VX-809 to ΔF508-CFTR and compared binding poses and affinities. Predicted binding affinities of PP2 and VX-809 by Autodock4 to the NBD1:ICL4 interface was −7.92 kcal/mol and −10.00 kcal/mol, respectively. In the case of the ATP binding site, the binding affinities for PP2 and VX-809 were −8.47 and −11.14 kcal/mol, respectively. These results suggest that PP2 might bind to the ATP binding site. The predicted binding pose of VRT-325 and PP2 is illustrated in Fig. 4D. Further, the heterobicyclic aromatic rings of PP2 positioned at the same place in the binding pocket, supporting our hypothesis that PP2 binds to the ATP site. To further compare the binding affinity of PP2 to the ATP binding site, we docked PP2 with four known binders (Fyn, Hck, Lck, and Src). Interestingly, the predicted binding affinities between PP2 and the four kinases were weaker than for PP2 and ATP binding site of CFTR. For instance, although the Lck protein is co-crystalized with PP2 (1QPE), its binding affinity was ∼1.8 kcal/mol lower than the ATP binding site of CFTR. This result strongly suggests that PP2 binds to the ATP binding site of CFTR.

### Characterization of PP2 as candidate therapeutic for CF

To gain insight into the mechanism of ΔF508-CFTR correction by PP2 in the context of CF pathophysiology, we analyzed the transcriptional profile of PP2 (from both LINCS and CF RNASeq data) along with CF DEGs.

We first analyzed the top upregulated and downregulated genes from PP2-treated cells (LINCS L1000 data) and DEGs from CF RE. Specifically, using our compiled CF- and CFTR-relevant biological processes, pathways, [33], and protein interactions (WT-CFTR and ΔF508-CFTR) [32], we performed functional enrichment analysis of these gene sets. PP2 upregulated genes including *TSPAN13, PRCP, RHOBTB3, TPD52L1, NRIP1, XBP1, SPDEF, MUC5B, HOXA5*, and *AGR2*, which were enriched in CF-related processes such as goblet cell differentiation and SPEDF induced genes. PP2 downregulated genes such as *PSMB1, NPLOC4, DYNLT1, DYNC1LI1, PSMB7, GNAI3, TUBB6, PPP2CB, TUBA4B, TUBB4B*, and *ADCY1*, which were enriched in CF-related processes including regulation of degradation of WT and DF508 CFTR and regulation of CFTR activity (normal and CF) (Fig. 5). Genes involved in CFTR proteostasis were enriched in upregulated DEGs in CFRE and top downregulated genes by PP2. These results suggest that PP2’s correction of CFTR might be by modulating degradation of mutant ΔF508-CFTR.

**Fig. 5.**
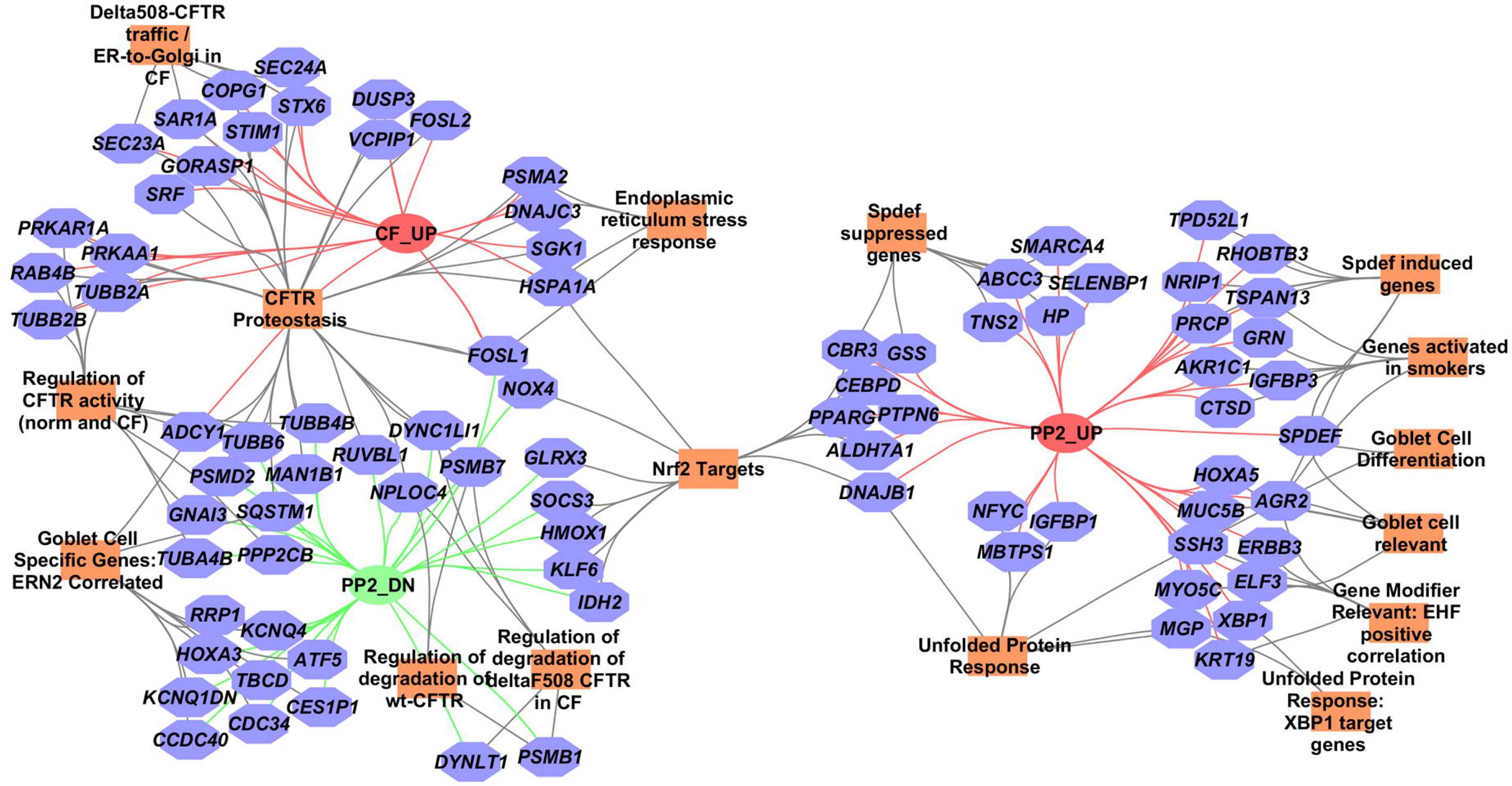
CF pathway enrichment network for genes that are reciprocally connected in CF and PP2 treatment. Rectangular orange nodes represent CF-related pathways; octagonal nodes represent differentially expressed genes either in CFRE or PP2-treated cells. Green edges represent downregulation; red edges represent upregulation.

Second, to gain context-specific insight into PP2 function, we performed RNA sequencing on ΔF508/ΔF508 *CFTR* and normal primary human bronchial epithelial cells treated with DMSO or PP2 (2 µM, 48 h). 119 genes were differentially expressed in PP2-treated bronchial epithelial cells compared with vehicle control (Fig. 6A). PP2 DEGs showed the highest positive connectivity with src-kinase inhibitors and the highest negative connectivity with tubulin inhibitors. Notably, PP2 partially reversed abnormal gene expression caused by F508del CFTR mutation in bronchial epithelial cells (Fig. 6B and 6C; Table 1, Additional file 5). These genes included matrix metalloprotease such as *MMP-1, MMP-9, MMP-10, MMP-12*, and *MMP-13*, which were downregulated by PP2. Among them, MMP-9 and MMP-12 had increased activity in CF patients [34, 35], and increased MMP-9 activity was negatively correlated with lung function [36]. PP2 also downregulated genes involved in CF-related pathway macrophage activation including *INHBA, IL1A*, and *FN1*. This is important because leukocyte inflammation in the airway is associated with increased CF severity in patients [37]. On the other hand, PP2 upregulated and restored expression of *CFI, C1R*, and *C1S*, which are involved in classical complement activation. This is potentially therapeutic since complement activation helps contain bacterial infection, which is an important contributing factor for CF pathogenesis in the lung [38, 39]. Taken together, these results suggest that PP2 could potentially protect the airway from inflammation and proteolytic damage in CF patients in addition to correcting the fundamental defect of mutant CFTR in CF (Fig. 7).

**Fig. 6:**
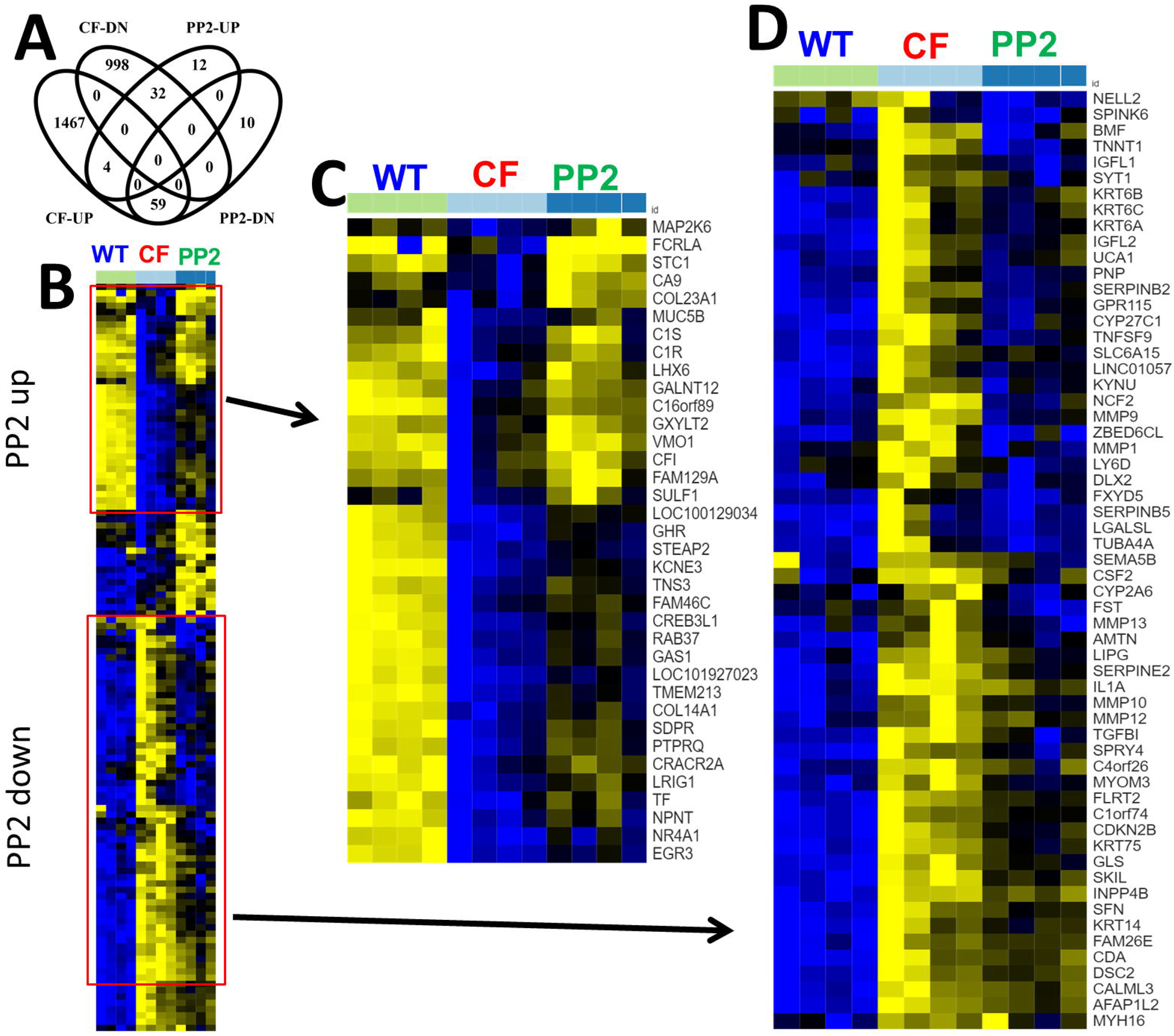
PP2 reversed the expression profiles of genes dysregulated in F508del-CFTR bronchial epithelial cells. Expression of 117 differentially expressed genes in bronchial epithelial cells with WT CFTR (green), F508del-CFTR (light blue), and F508del-CFTR treated with 2 µM PP2 (dark blue) is shown. Each row represents a gene, and each column represents a sample. Colors were mapped based on Reads Per Kilobase of transcript per Million mapped reads (RPKM) values, where yellow corresponds to high expression and blue corresponds to low expression.

**Fig. 7:**
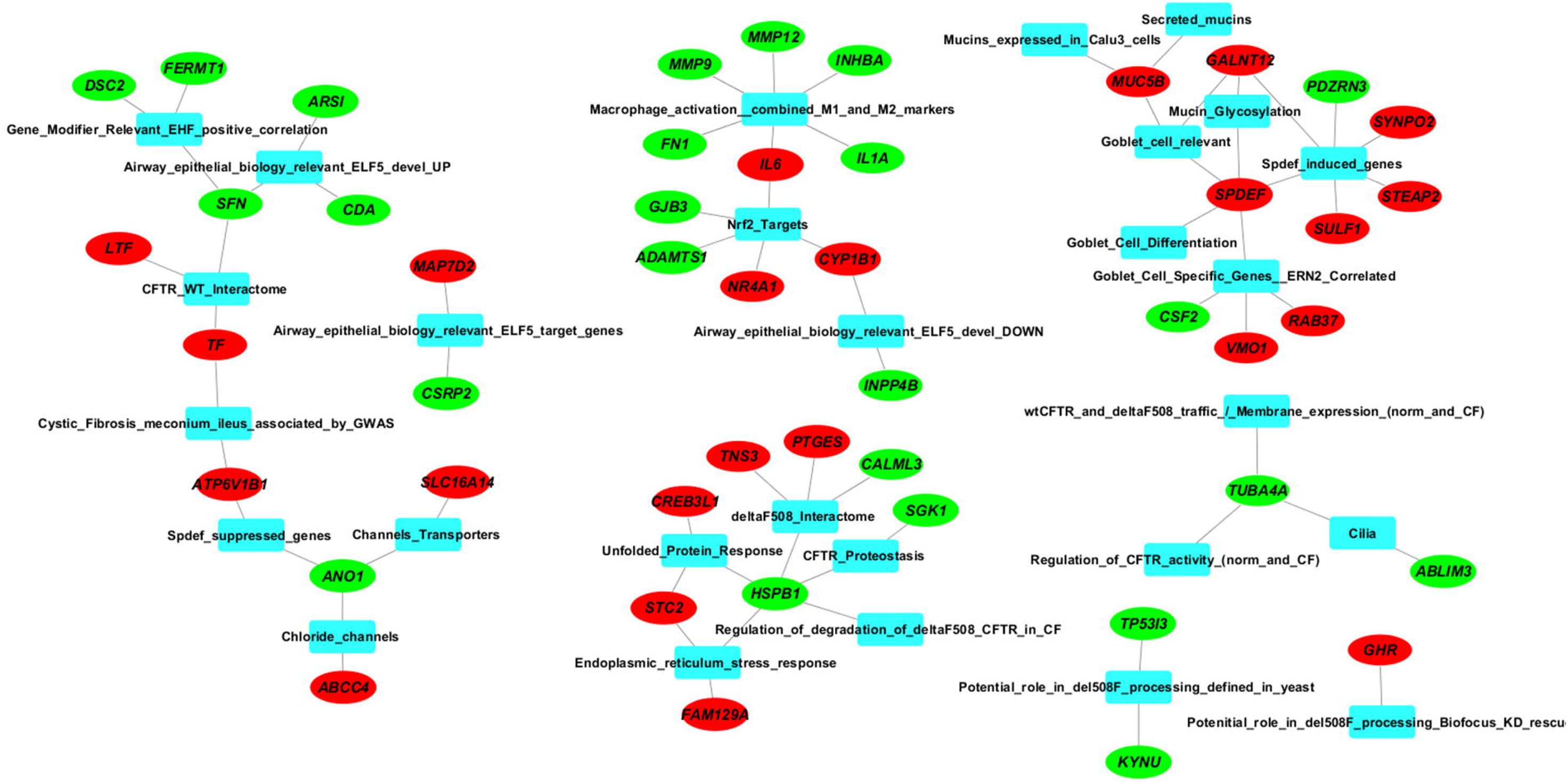
Functional enrichment network of differentially expressed genes following treatment with PP2 (2 µM; 48 h) in human CFBE (ΔF508/ΔF508-*CFTR*)

**Table 1:**
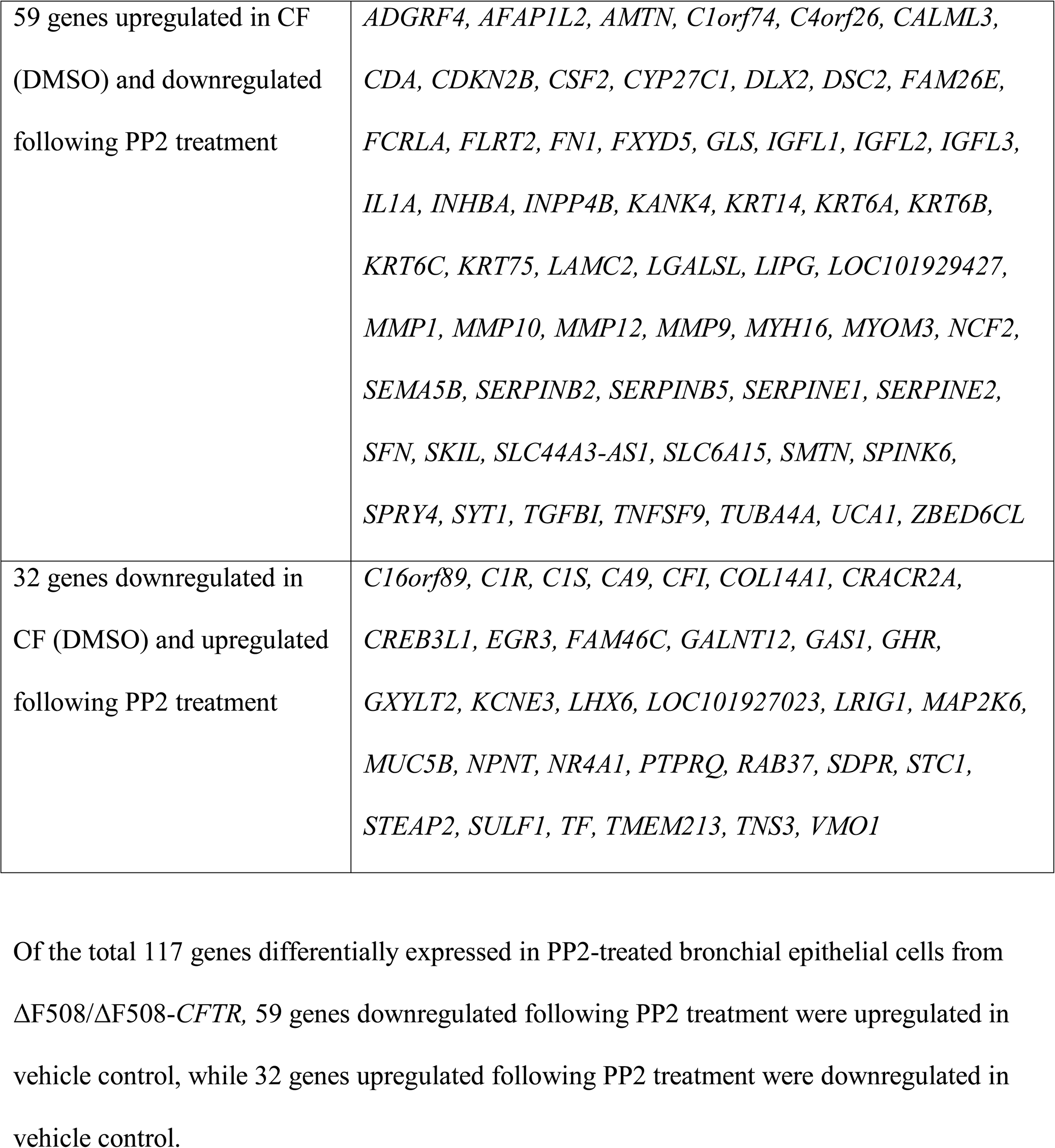
Comparison of genes differentially expressed in DMSO-treated ΔF508/ΔF508-*CFTR* primary human bronchial epithelial cells with those following treatment with PP2 (2 µM, 48 h).

## Discussion

In this study, we described a CF-specific systems biology-guided unbiased computational compound screening to identify and prioritize novel small molecules that could potentially rescue ΔF508-CFTR function. Using enteroids generated from the ileum of ΔF508-CFTR homozygous mice and from rectal biopsy of CF patients (ΔF508-CFTR homozygous) and *I_sc_* analysis of human bronchial epithelial cells harvested from homozygous ΔF508-CFTR transplant patients (extensive functional validation), we validated our computationally predicted and prioritized small-molecule candidates. Recent studies have shown the utility of using CF patient-derived rectal organoids for drug screening [40].

Based on our findings, we reported a novel corrector (PP2, a src-kinase inhibitor) of the ΔF508-CFTR defect. Connectivity mapping of two other gene expression data sets from CF (human CF bronchial epithelia [41] and CF Pigs [42]) with LINCS signatures also showed PP2 among the top hits (Additional file 6). Our proof-of-principle studies also demonstrated src-kinase inhibitors as a new class of compounds that can be used for rescuing ΔF508-CFTR.

While we observed synergy with VX-809, the potential side effects of PP2 in combination with VX-809 (or VX-661) are difficult to predict. Further, we noted that PP2, like most other src-kinase inhibitors, might be mechanistically associated with drug-induced side effects. Thus, thorough toxicity tests in animal models followed by clinical studies of PP2 and the combinations (PP2 and VX-809/VX-661) would have to be performed.

Although we tested 24 compounds from a total of 184 candidates, we believe that additional CF candidate therapeutics can be identified from the remaining nontested compounds. For instance, among the remaining 160 compounds, we further prioritized 30 compounds that are highly similar to both PP2 and LY-294002 as CF candidate therapeutics (Additional file 7).

Although results from our in vitro analysis and computational docking (Figs. 4B and 4D) suggest potential direct action of PP2 on mutant CFTR, additional in vitro and in vivo analysis are warranted. For instance, is the PP2-induced mutant CFTR correction dependent on the canonical SFK (src family kinase) inhibition? SFKs have important roles in biological processes altered in CF such as apoptosis, inflammatory response, autophagy and mucin production, although the exact relation between SFK and these processes in the context of CF remain largely unknown [43]. A recent study showed that CFTR deficiency leads to SFK self-activation and intensified inflammatory response in mouse cholangiocytes exposed to endotoxin or LPS, and inhibition of SFK with PP2 decreased inflammation [44]. This is in consistence with our results wherein we found that PP2 downregulated genes in human CFBE showed an enrichment (*P*<0.05) for macrophage activation markers (Additional file 5) such as *INHBA*, *IL1A*, *MMP12*, *MMP9*, and *FN1*, suggesting that PP2 may potentially ameliorate SFK-triggered inflammatory cascades.

Recurrent *Pseudomonas aeruginosa* (PsA) infection is a major contributor to CF pathology [45]. Previous studies [46] have shown that Lyn, an SFK member is critical for *PsA* internalization into lung cells and that blocking the activity of Lyn with PP2 prevented *PsA* internalization. It has also been shown that the NLRP3 inflammasome is mediated by SFK activity [47] and that NLRP3 activation exacerbates the *PsA*-driven inflammatory response in CF [48] and NLRP3 inflammasomes are potential targets to limit the microbial colonization in CF [49]. Additionally, it has been reported that *PsA* infection induces the expression of MMPs (MMP12 and MMP13), which further exacerbates chronic lung infection and inflammation [50, 51]. In our study, we found PP2 downregulated MMPs and *NLRP3* in CFBE treated with PP2 (Additional file 5), suggesting PP2 could also be therapeutic in CF though inhibition of PsA infection.

## Conclusions

In addition to novel candidate drugs to target mutated CFTR, novel anti-inflammatory and anti-infective drugs that address secondary disease pathology in CF are needed to lessen morbidity, prolong survival, and improve quality of life [52, 53]. Thus, although additional studies are required, our results suggest that the SFK-inhibition (e.g., PP2) may represent a novel paradigm of multi-action therapeutics – corrector, anti-inflammatory, and anti-infective – in CF. Interestingly, while this study was underway, a second study from Strazzabosco lab reported [54] that the combination of correctors and PP2 in ΔF508 cholangiocytes significantly increased the amount of the Band C of ΔF508-CFTR.

## Abbreviations

CF: Cystic fibrosis
CFTR: CF transmembrane conductance regulator
LINCS: Library of Integrated Network-based Cellular Signatures
FIS: Forskolin-induced-swelling
CFRE: CF rectal epithelia
*I*_sc_: short circuit currents
DEG: differentially expressed gene
GSEA: gene set enrichment analysis
SEA: singular enrichment analysis
*PsA*: *Pseudomonas aeruginosa*
PP2: 1-tert-Butyl-3-(4-chlorophenyl)-1H-pyrazolo[3,4-d]pyrimidin-4-amine

## Declarations

### Ethics approval and consent to participate

*Human studies*: Patient-derived organoid assay studies for research were approved by CCHMC IRB under 2011-2616.

*Mice studies*: All procedures in mice were performed in compliance with institutional guidelines and were approved by CCHMC’s IACUC.

## Consent for publication

Not applicable.

## Availability of data and material

All data generated or analyzed during this study are included in this published article and its supplementary files.

## Authors’ contributions

YW, KA, APN, and AGJ conceived the project, designed experiments, analyzed data, and wrote the manuscript. YW and JC helped with the data analysis. FY helped with the experiments. WH and DK performed the docking studies. All the authors have read and approved the final manuscript.

## Acknowledgement

The authors thank David L. Armbruster for editing the manuscript. The authors would like to thank Drs. Jeffrey Whitsett and Bruce Aronow for their support.

## Additional files

Additional file 1: All differentially expressed genes in the CFRE dataset (GSE15568).

Additional file 2: CF-related/relevant pathways and gene sets.

Additional file 3: List of 184 compounds identified by LINCS L1000 compound screening.

Additional file 4: 24 compounds shortlisted for enteroid assay. This file also includes the raw data related to FIS assay for the 24 compounds along with the WT, VX809, and VX661.

Additional file 5: All differentially expressed genes in human CFBE cells treated with PP2.

Additional file 6: Comparison of compounds prioritized with additional data sets

Additional file 7: List of 30 compounds that are highly similar to both PP2 and LY-294002 as additional CF candidate therapeutics

## References

1. Dekkers JF, van der Ent CK, Beekman JM: Novel opportunities for CFTR-targeting drug development using organoids. Rare Dis 2013, 1:e27112.

2. Cholon DM, Quinney NL, Fulcher ML, Esther CR, Jr., Das J, Dokholyan NV, Randell SH, Boucher RC, Gentzsch M: Potentiator ivacaftor abrogates pharmacological correction of DeltaF508 CFTR in cystic fibrosis. Sci Transl Med 2014, 6:246ra296.

3. Veit G, Avramescu RG, Perdomo D, Phuan PW, Bagdany M, Apaja PM, Borot F, Szollosi D, Wu YS, Finkbeiner WE, et al: Some gating potentiators, including VX-770, diminish DeltaF508-CFTR functional expression. Sci Transl Med 2014, 6:246ra297.

4. Phuan PW, Veit G, Tan JA, Finkbeiner WE, Lukacs GL, Verkman AS: Potentiators of Defective DeltaF508-CFTR Gating that Do Not Interfere with Corrector Action. Mol Pharmacol 2015, 88:791–799.

5. Clancy JP: CFTR potentiators: not an open and shut case. Sci Transl Med 2014, 6:246fs227.

6. Boyle MP, Bell SC, Konstan MW, McColley SA, Rowe SM, Rietschel E, Huang X, Waltz D, Patel NR, Rodman D, group VXs: A CFTR corrector (lumacaftor) and a CFTR potentiator (ivacaftor) for treatment of patients with cystic fibrosis who have a phe508del CFTR mutation: a phase 2 randomised controlled trial. Lancet Respir Med 2014, 2:527–538.

7. Dudley JT, Sirota M, Shenoy M, Pai RK, Roedder S, Chiang AP, Morgan AA, Sarwal MM, Pasricha PJ, Butte AJ: Computational repositioning of the anticonvulsant topiramate for inflammatory bowel disease. Sci Transl Med 2011, 3:96ra76.

8. Sirota M, Dudley JT, Kim J, Chiang AP, Morgan AA, Sweet-Cordero A, Sage J, Butte AJ: Discovery and preclinical validation of drug indications using compendia of public gene expression data. Sci Transl Med 2011, 3:96ra77.

9. Iorio F, Bosotti R, Scacheri E, Belcastro V, Mithbaokar P, Ferriero R, Murino L, Tagliaferri R, Brunetti-Pierri N, Isacchi A, di Bernardo D: Discovery of drug mode of action and drug repositioning from transcriptional responses. Proc Natl Acad Sci U S A 2010, 107:14621–14626.

10. Iorio F, Isacchi A, di Bernardo D, Brunetti-Pierri N: Identification of small molecules enhancing autophagic function from drug network analysis. Autophagy 2010, 6:1204–1205.

11. Qu XA, Gudivada RC, Jegga AG, Neumann EK, Aronow BJ: Inferring novel disease indications for known drugs by semantically linking drug action and disease mechanism relationships. BMC Bioinformatics 2009, 10.

12. Qu XA, Freudenberg JM, Sanseau P, Rajpal DK: Integrative clinical transcriptomics analyses for new therapeutic intervention strategies: a psoriasis case study. Drug Discov Today 2014, 19:1364–1371.

13. Ramachandran S, Osterhaus SR, Karp PH, Welsh MJ, McCray PB, Jr.: A genomic signature approach to rescue DeltaF508-cystic fibrosis transmembrane conductance regulator biosynthesis and function. Am J Respir Cell Mol Biol 2014, 51:354–362.

14. Lamb J, Crawford ED, Peck D, Modell JW, Blat IC, Wrobel MJ, Lerner J, Brunet JP, Subramanian A, Ross KN, et al: The Connectivity Map: using gene-expression signatures to connect small molecules, genes, and disease. Science 2006, 313:1929–1935.

15. Cheng J, Yang L, Kumar V, Agarwal P: Systematic evaluation of connectivity map for disease indications. Genome Med 2014, 6.

16. Hu G, Agarwal P: Human disease-drug network based on genomic expression profiles. PLoS One 2009, 4:e6536.

17. Qu XA, Rajpal DK: Applications of Connectivity Map in drug discovery and development. Drug Discov Today 2012, 17:1289–1298.

18. Stanke F, van Barneveld A, Hedtfeld S, Wolfl S, Becker T, Tummler B: The CF-modifying gene EHF promotes p.Phe508del-CFTR residual function by altering protein glycosylation and trafficking in epithelial cells. Eur J Hum Genet 2014, 22:660–666.

19. Library of Integrated Cellular Signatures (LINCS) [http://www.lincsproject.org/]

20. Subramanian A, Narayan R, Corsello SM, Peck DD, Natoli TE, Lu X, Gould J, Davis JF, Tubelli AA, Asiedu JK, et al: A Next Generation Connectivity Map: L1000 Platform and the First 1,000,000 Profiles. Cell 2017, 171:1437–1452 e1417.

21. Li C, Dandridge KS, Di A, Marrs KL, Harris EL, Roy K, Jackson JS, Makarova NV, Fujiwara Y, Farrar PL, et al: Lysophosphatidic acid inhibits cholera toxin-induced secretory diarrhea through CFTR-dependent protein interactions. J Exp Med 2005, 202:975–986.

22. Li C, Krishnamurthy PC, Penmatsa H, Marrs KL, Wang XQ, Zaccolo M, Jalink K, Li M, Nelson DJ, Schuetz JD, Naren AP: Spatiotemporal coupling of cAMP transporter to CFTR chloride channel function in the gut epithelia. Cell 2007, 131:940–951.

23. Barrett T, Troup DB, Wilhite SE, Ledoux P, Rudnev D, Evangelista C, Kim IF, Soboleva A, Tomashevsky M, Edgar R: NCBI GEO: mining tens of millions of expression profiles--database and tools update. Nucleic Acids Res 2007, 35:D760–765.

24. Ritchie ME, Phipson B, Wu D, Hu YF, Law CW, Shi W, Smyth GK: limma powers differential expression analyses for RNA-sequencing and microarray studies. Nucleic Acids Research 2015, 43.

25. Ooms J: The jsonlite Package: A Practical and Consistent Mapping Between JSON Data and R Objects. In ArXiv e-prints, vol. 1403; 2014.

26. Dalton J, Kalid O, Schushan M, Ben-Tal N, Villa-Freixa J: New model of cystic fibrosis transmembrane conductance regulator proposes active channel-like conformation. J Chem Inf Model 2012, 52:1842–1853.

27. Sali A, Blundell TL: Comparative protein modelling by satisfaction of spatial restraints. J Mol Biol 1993, 234:779–815.

28. Kim S, Thiessen PA, Bolton EE, Chen J, Fu G, Gindulyte A, Han L, He J, He S, Shoemaker BA, et al: PubChem Substance and Compound databases. Nucleic Acids Res 2016, 44:D1202–1213.

29. Morris GM, Huey R, Lindstrom W, Sanner MF, Belew RK, Goodsell DS, Olson AJ: AutoDock4 and AutoDockTools4: Automated docking with selective receptor flexibility. J Comput Chem 2009, 30:2785–2791.

30. Moon C, Zhang W, Ren A, Arora K, Sinha C, Yarlagadda S, Woodrooffe K, Schuetz JD, Valasani KR, de Jonge HR, et al: Compartmentalized accumulation of cAMP near complexes of multidrug resistance protein 4 (MRP4) and cystic fibrosis transmembrane conductance regulator (CFTR) contributes to drug-induced diarrhea. J Biol Chem 2015, 290:11246–11257.

31. Dekkers JF, Wiegerinck CL, de Jonge HR, Bronsveld I, Janssens HM, de Winter-de Groot KM, Brandsma AM, de Jong NW, Bijvelds MJ, Scholte BJ, et al: A functional CFTR assay using primary cystic fibrosis intestinal organoids. Nat Med 2013, 19:939–945.

32. Pankow S, Bamberger C, Calzolari D, Martinez-Bartolome S, Lavallee-Adam M, Balch WE, Yates JR, 3rd: F508 CFTR interactome remodelling promotes rescue of cystic fibrosis. Nature 2015, 528:510–516.

33. O’Neal WK, Gallins P, Pace RG, Dang H, Wolf WE, Jones LC, Guo X, Zhou YH, Madar V, Huang J, et al: Gene expression in transformed lymphocytes reveals variation in endomembrane and HLA pathways modifying cystic fibrosis pulmonary phenotypes. Am J Hum Genet 2015, 96:318–328.

34. Gaggar A, Li Y, Weathington N, Winkler M, Kong M, Jackson P, Blalock JE, Clancy JP: Matrix metalloprotease-9 dysregulation in lower airway secretions of cystic fibrosis patients. Am J Physiol Lung Cell Mol Physiol 2007, 293:L96–L104.

35. Ratjen F, Hartog CM, Paul K, Wermelt J, Braun J: Matrix metalloproteases in BAL fluid of patients with cystic fibrosis and their modulation by treatment with dornase alpha. Thorax 2002, 57:930–934.

36. Sagel SD, Kapsner RK, Osberg I: Induced sputum matrix metalloproteinase-9 correlates with lung function and airway inflammation in children with cystic fibrosis. Pediatr Pulmonol 2005, 39:224–232.

37. Gaggar A, Hector A, Bratcher PE, Mall MA, Griese M, Hartl D: The role of matrix metalloproteinases in cystic fibrosis lung disease. Eur Respir J 2011, 38:721–727.

38. Dunkelberger JR, Song WC: Complement and its role in innate and adaptive immune responses. Cell Res 2010, 20:34–50.

39. Nichols DP, Chmiel JF: Inflammation and its genesis in cystic fibrosis. Pediatr Pulmonol 2015, 50 Suppl 40:S39–56.

40. Dekkers JF, Berkers G, Kruisselbrink E, Vonk A, de Jonge HR, Janssens HM, Bronsveld I, van de Graaf EA, Nieuwenhuis EE, Houwen RH, et al: Characterizing responses to CFTR-modulating drugs using rectal organoids derived from subjects with cystic fibrosis. Sci Transl Med 2016, 8:344ra384.

41. Ogilvie V, Passmore M, Hyndman L, Jones L, Stevenson B, Wilson A, Davidson H, Kitchen RR, Gray RD, Shah P, et al: Differential global gene expression in cystic fibrosis nasal and bronchial epithelium. Genomics 2011, 98:327–336.

42. Stoltz DA, Meyerholz DK, Pezzulo AA, Ramachandran S, Rogan MP, Davis GJ, Hanfland RA, Wohlford-Lenane C, Dohrn CL, Bartlett JA, et al: Cystic fibrosis pigs develop lung disease and exhibit defective bacterial eradication at birth. Sci Transl Med 2010, 2:29ra31.

43. Massip Copiz MM, Santa Coloma TA: c-Src and its role in cystic fibrosis. Eur J Cell Biol 2016, 95:401–413.

44. Fiorotto R, Villani A, Kourtidis A, Scirpo R, Amenduni M, Geibel PJ, Cadamuro M, Spirli C, Anastasiadis PZ, Strazzabosco M: The cystic fibrosis transmembrane conductance regulator controls biliary epithelial inflammation and permeability by regulating Src tyrosine kinase activity. Hepatology 2016, 64:2118–2134.

45. Bhagirath AY, Li Y, Somayajula D, Dadashi M, Badr S, Duan K: Cystic fibrosis lung environment and Pseudomonas aeruginosa infection. BMC Pulm Med 2016, 16:174.

46. Kannan S, Audet A, Knittel J, Mullegama S, Gao GF, Wu M: Src kinase Lyn is crucial for Pseudomonas aeruginosa internalization into lung cells. Eur J Immunol 2006, 36:1739–1752.

47. Guo W, Ye P, Yu H, Liu Z, Yang P, Hunter N: CD24 activates the NLRP3 inflammasome through c-Src kinase activity in a model of the lining epithelium of inflamed periodontal tissues. Immun Inflamm Dis 2014, 2:239–253.

48. Rimessi A, Bezzerri V, Patergnani S, Marchi S, Cabrini G, Pinton P: Mitochondrial Ca2+-dependent NLRP3 activation exacerbates the Pseudomonas aeruginosa-driven inflammatory response in cystic fibrosis. Nat Commun 2015, 6:6201.

49. Iannitti RG, Napolioni V, Oikonomou V, De Luca A, Galosi C, Pariano M, Massi-Benedetti C, Borghi M, Puccetti M, Lucidi V, et al: IL-1 receptor antagonist ameliorates inflammasome-dependent inflammation in murine and human cystic fibrosis. Nat Commun 2016, 7:10791.

50. Park JW, Kim YJ, Shin IS, Kwon OK, Hong JM, Shin NR, Oh SR, Ha UH, Kim JH, Ahn KS: Type III Secretion System of Pseudomonas aeruginosa Affects Matrix Metalloproteinase 12 (MMP-12) and MMP-13 Expression via Nuclear Factor κB Signaling in Human Carcinoma Epithelial Cells and a Pneumonia Mouse Model. J Infect Dis 2016, 214:962–969.

51. Park JW, Shin IS, Ha UH, Oh SR, Kim JH, Ahn KS: Pathophysiological changes induced by Pseudomonas aeruginosa infection are involved in MMP-12 and MMP-13 upregulation in human carcinoma epithelial cells and a pneumonia mouse model. Infect Immun 2015, 83:4791–4799.

52. Chmiel JF, Konstan MW, Elborn JS: Antibiotic and anti-inflammatory therapies for cystic fibrosis. Cold Spring Harb Perspect Med 2013, 3:a009779.

53. Nichols DP, Konstan MW, Chmiel JF: Anti-inflammatory therapies for cystic fibrosis-related lung disease. Clin Rev Allergy Immunol 2008, 35:135–153.

54. Fiorotto R, Amenduni M, Mariotti V, Fabris L, Spirli C, Strazzabosco M: Src kinase inhibition reduces inflammatory and cytoskeletal changes in ΔF508 human cholangiocytes and improves CFTR correctors efficacy. Hepatology 2017.

